# Decreased Colonic Activin Receptor-Like Kinase 1 Disrupts Epithelial Barrier integrity and is associated with a poor clinical outcome in Crohn’s disease

**DOI:** 10.1101/2020.02.21.960070

**Authors:** Takahiko Toyonaga, Benjamin P. Keith, Jasmine B. Barrow, Matthew S. Schaner, Elisabeth A. Wolber, Caroline Beasley, Jennifer Huling, Yuli Wang, Nancy L. Allbritton, Nicole Chaumont, Timothy S. Sadiq, Mark J. Koruda, Reza Rahbar, Terrence S. Furey, Praveen Sethupathy, Shehzad Z. Sheikh

## Abstract

**Objective:** Intestinal epithelial cell (IEC) barrier dysfunction is critical to the development of Crohn’s disease (CD). However, the mechanism is understudied. We recently reported increased microRNA-31-5p (miR-31-5p) expression in colonic IECs of CD patients, but downstream targets are unknown.

**Design:** MiR-31-5p target genes were identified by integrative analysis of RNA- and small RNA-sequencing data from colonic mucosa and confirmed by qPCR in colonic IECs. Functional characterization of Activin Receptor-Like Kinase 1 (ACVRL1 or ALK1) in IECs was performed *ex vivo* using 2 dimensional-cultured human primary colonic IECs. The impact of altered colonic ALK1 signaling in CD for the risk of surgery and endoscopic relapse was evaluated by a multivariate regression analysis and a Kaplan-Meier estimator.

**Results:** *ALK1* was identified as a target of miR-31-5p in colonic IECs of CD patients and confirmed using a 3’-UTR reporter assay. Activation of ALK1 restricted the proliferation of colonic IECs in an EdU proliferation assay and down-regulated the expression of stemness-related genes. Activated ALK1 signaling directed the fate of colonic IEC differentiation toward colonocytes. Down-regulated ALK1 signaling was associated with increased stemness and decreased colonocyte-specific marker expression in colonic IECs of CD patients compared to healthy controls. Activation of ALK1 enhanced epithelial barrier integrity in a trans-epithelial electrical resistance permeability assay. Lower colonic *ALK1* expression was identified as an independent risk factor for surgery and associated with a higher risk of endoscopic relapse in CD patients.

**Conclusion:** Decreased colonic ALK1 disrupted colonic IEC barrier integrity and associated with deteriorated clinical outcomes in CD patients.

## Introduction

Chronic intestinal inflammation in Crohn’s disease (CD) is caused by an aberrant interaction between the mucosal immune system and luminal antigens, often leading to a compromised IEC barrier, in genetically predisposed individuals^1^. There is an increasing appreciation for the importance of variable cellular processes in intestinal epithelial cells (IECs) that contribute to the loss of barrier capacity and the development of CD^2, 3^. We^4-6^ and others^7^ have shown that alterations in IEC gene expression are associated with CD, but the underlying mechanisms driving IEC barrier defects remain unresolved. Current management of CD is largely directed at aberrant immune responses while CD therapy targeting IEC barrier defects remain elusive^8^.

In contrast to the small intestine where Paneth cells provide additional support for epithelial barrier integrity in part by secreting antimicrobial peptides, the colonic IEC barrier is largely maintained by tight cell-to-cell connections between colonocytes and by the mucus secreted from goblet cells^9^. Colonocytes are also enriched with junctional proteins^10^ that serve as a barrier to luminal contents. Defects in the colonocyte barrier can increase paracellular permeability, which has been shown to lead to intestinal inflammation in several rodent models^11, 12^. Mechanisms controlling human colonocyte differentiation and survival are still understudied, especially in CD patients.

We recently reported aberrantly elevated expression of microRNA-31-5p (miR-31-5p) in colonic IECs of CD patients compared to non-inflammatory bowel disease (NIBD) controls^4^. miRNAs contribute to the control of various biological processes, including proliferation and differentiation^13^, by post-transcriptionally regulating target gene expression^14^. We hypothesized that miR-31-5p might contribute to the pathophysiology of CD through regulation of key genes that drive colonic IEC proliferation and differentiation. In this study, we identified and validated a novel miR-31-5p target gene, Activin Receptor-Like Kinase 1 (*ACVRL1* or *ALK1*), in human colonic IECs. The functions of ALK1 in the colon have not previously been investigated. We examined the functional impact of ALK1 signaling on IEC biology, notably colonocyte differentiation and barrier integrity, using human primary IECs. We also examined the clinical impact of altered colonic ALK1 in patients with CD and demonstrated an association of decreased colonic *ALK1* with increased risks of surgery and endoscopic relapse.

## Materials and Methods

### Subjects, Samples, and Clinical Information

Colonic mucosa was obtained from surgically resected colon or endoscopically taken biopsies from patients with established diagnosis of CD and NIBD healthy controls between April 2012 and November 2019. Cross-sectional clinical data was collected at the time of sampling. All samples were collected from disease-unaffected regions without macroscopic inflammation. All biopsy samples were collected from ascending colon. Additional information for surgical samples is summarized in **Supplementary Table 1**. Endoscopic disease activity was evaluated by Simple Endoscopic Score for Crohn’s Disease (SES-CD)^15^ in CD patients. Information required for scoring was retrospectively extracted from endoscopy reports. Endoscopic remission was defined as SES-CD less than 4^16^. In patients with endoscopic remission, endoscopic disease activity was prospectively followed up to 72 weeks after sampling to examine endoscopic relapse, defined as SES-CD ≧4.

### Isolation of Colonic Crypts and Primary Culture of Human Intestinal Epithelial Cells

Isolation of colonic crypts and subsequent human primary IEC culture were performed as we reported previously^17, 18^. In brief, colonic mucosa was incubated in the isolation buffer^17, 18^ for 75 minutes at room temperature. The tissue was rinsed with Phosphate Buffered Saline (PBS) and vigorously shaken by hand to release the crypts. The released crypts were placed into culture immediately or suspended in expansion medium (EM)^18^ containing 20% FBS and 10% dimethyl sulfoxide (DMSO) to store at −80°C until use.

For two-dimensional (2D) culture, isolated crypts were directly spread on 48-well plates coated with collagen hydrogel^17^. Cells were cultured in EM with 10mM Y-27632 (Selleckchem, TX, USA) during the initial 48 hours. The medium was changed every 48 hours. Every 6 days, cells were dissociated in collagenase type IV followed by Accutase (Stemcell Technologies, BC, Canada) and the fragmented cells were passaged to a new collagen-coated plate.

In several experiments, cells were expanded in EM for 4 days and subsequently stimulated with 10 ng/ml bone morphogenetic protein (BMP) 9 (BioLegend, CA, USA) and 100 ng/ml ALK1-Fc chimera protein (R&D Systems, MN, USA), a soluble chimeric protein consisting of the extracellular part of ALK1 fused to a Fc fragment, which inhibits BMP9-ALK1 interaction and signaling^19^.

### Epithelial Permeability Assay

Epithelial permeability was evaluated in 2D-cultured human primary IEC monolayers as we reported previously^20^. Primary IECs cultured on 48-well plates were passaged to 12-well cell culture inserts (transparent polyethylene terephthalate membrane, membrane pore size 0.4 um, effective cell culture area 0.9 cm^2^; Corning, NY, USA) coated with collagen scaffolds with a gradient of cross-linking^21^ at a ratio of 1:1. Cells were expanded in EM for 4 days and stimulated with 10 ng/ml BMP9 and 100 ng/ml ALK1-Fc chimera protein in EM or cultured in differentiation medium (DM) for additional 2 days. In contrast to EM, DM lack exogeneous Wnt Family Member 3A (WNT3A), R-SPONDIN 3, and NOGGIN produced from L-WRN cells^18^ and SB202190 (Selleckchem), a selective p38 mitogen-activated protein kinase inhibitor. The resistance between the upper and lower surfaces of IEC monolayers was measured before (Day 4) and after 2 days of stimulation (Day 6) using an EVOM2 epithelial Volt/Ohm meter (World Precision Instruments, FL, USA). The resistance of IEC monolayers was corrected by subtracting the resistance of scaffolds without cells and normalized by multiplying the effective scaffold surface area to provide a trans-epithelial electrical resistance (TEER) in the unit of Ω cm^2^. Changes in TEER between Day4 and Day6 were evaluated by calculating delta TEER% as following: ΔTEER% =100- [{(TEER_Day4_-TEER_Day6_)/TEER_Day4_} x 100]^20^.

### 3’UTR Reporter Assay and Site-directed Mutagenesis

Plasmid DNA was extracted from a miRNA 3’UTR target clone for ALK1 (HmiT022834-MT06, GeneCopoeia, MD, USA) using Qiagen Plasmid Midi Kit (Qiagen, MD, USA). This plasmid DNA was then transfected in HEK293T cells with or without miRNA mimics, double-stranded oligonucleotides designed to mimic the function of endogenous mature miRNAs, for hsa-miR-31-5p, hsa-miR-122a-5p, or hsa-miR-215-5p, or negative control mimics (Dharmacon, CO, USA). Cells transfected with plasmid DNA in the absence of miRNA mimics were defined as mock. For mutagenesis assays, putative miR-31-5p binding sequences were deleted in the plasmid DNA using PrimeSTAR Mutagenesis Basal Kit (TaKaRa Bio, CA, USA) according to the manufacturer’s instructions. Successful deletions were checked by Sanger DNA sequencing (Eurofins Genomics) and subsequently analyzed with Sequencher 5.4.6 (Gene Codes). Dual luciferase reporter assays were performed on Dual Luciferase Reporter Assay System (Promega, WI, USA) using the Luc-Pair Duo-Luciferase HS Assay Kit (GeneCopoeia) according to the manufacturer’s instructions.

### 5-Ethynyl-2-deoxyuridine assay

The 5-ethynyl-2-deoxyuridine (EdU)-based staining was performed on 2D-cultured primary IECs. Cells were sub-cultured from 48-well plates to 96-well plates at a ratio of 1:3, expanded in EM for 4 days followed by additional 2 days in EM with or without 10 ng/ml BMP9 and 100 ng/ml ALK1-Fc chimera protein or in DM. Cells were incubated with 10mM EdU for 3 hours at 37°C. DNA-incorporated EdU was detected by Sulfo-Cyanine5-azide (Lumiprobe, MD, USA) through a copper-catalyzed covalent reaction between an azide and alkyne^22^. Images were captured using a Nikon Eclipse TE2000-U inverted microscope (Nikon, Tokyo, Japan) and processed using NIS-Elements AR version 3.2 software. The EdU fluorescence area was measured by ImageJ software and normalized by the total cell area occupied by the Hoechst 33342 fluorescence as we described previously^18^.

### Statistical analysis

All numerical data in the figures are expressed as means ± standard deviation (SD) or standard error of the mean (SEM). Differences between two groups were analyzed by Wilcoxon test, Mann-Whitney test or Fisher’s exact test. Differences among three groups were analyzed by Kruskal-Wallis test or Friedman test followed by Dunn’s multiple comparison test. Trends between variables were analyzed by Jonckheere-Terpstra test.

Spearman’s correlation coefficient was used for evaluating correlations between two numerical variables. Correlations between two categorical variables were evaluated by chi-squared test with Cramer’s V to measure the strength of correlations. *P*-values less than 0.05 were considered significant. The Kaplan-Meier method was used to generate survival curves and differences between two groups were evaluated by log-rank test. The GraphPad Prism version 8.0 software (GraphPad Software, CA, USA) and R software version 3.5.2 were used for these data analyses. A binomial logistic regression analysis was also performed using R software.

### Ethical statement

The study was conducted in accordance with the Declaration of Helsinki and Good Clinical Practice. The study protocol was approved by the institutional review board at University of North Carolina at Chapel Hill (approval number: 19-0819 and 17-0236). All participants provided written informed consent prior to inclusion in the study. All participants are identified by number and not by name or any protected health information.

## Results

### ALK1 is a putative target of miR-31-5p in human colonic epithelial cells

We previously generated and studied the expression profiles of mRNAs^6^ and miRNAs^4^ in the uninflamed colonic mucosa of 21 adult CD patients and 11 NIBD controls revealing a unique subset of CD patients with especially high miR-31-5p expression^23^. Predicted miR-31-5p target genes (n=27) were significantly enriched among the genes downregulated in this group of “high-miR-31-5p” CD patients compared to NIBD controls^4^ (**Supplementary Table 2**). Correlation analysis in the high-miR-31-5p CD patients revealed that 4 of these genes were significantly inversely correlated with miR-31 (**Figure 1A**). The strongest two of these were *E2F Transcription Factor 2* (*E2F2*) and *Activin Receptor-Like Kinase 1* (*ALK1*) (r = −0.92 and −0.90; p<0.001 and p<0.001, respectively). We next examined the association of miR-31-5p with *E2F2* and *ALK1* specifically in colonic IECs isolated from CD patients and NIBD controls by qPCR (**Figure 1B**), which revealed a significant negative correlation between miR-31-5p and *ALK1* but not *E2F2* (*ALK1*, p=0.019; *E2F2*, p=0.982, Jonckheere-Terpstra test). Relative ALK1 protein expression was significantly inversely correlated with miR-31-5p expression (r = −0.59, p<0.01) and was remarkably decreased in the colonic mucosa of CD patients with higher miR-31-5p expression compared to NIBD controls (**Figure 1C**). This suggests that *ALK1* is a putative target of miR-31-5p in human colonic IECs.

**Fig. 1.**
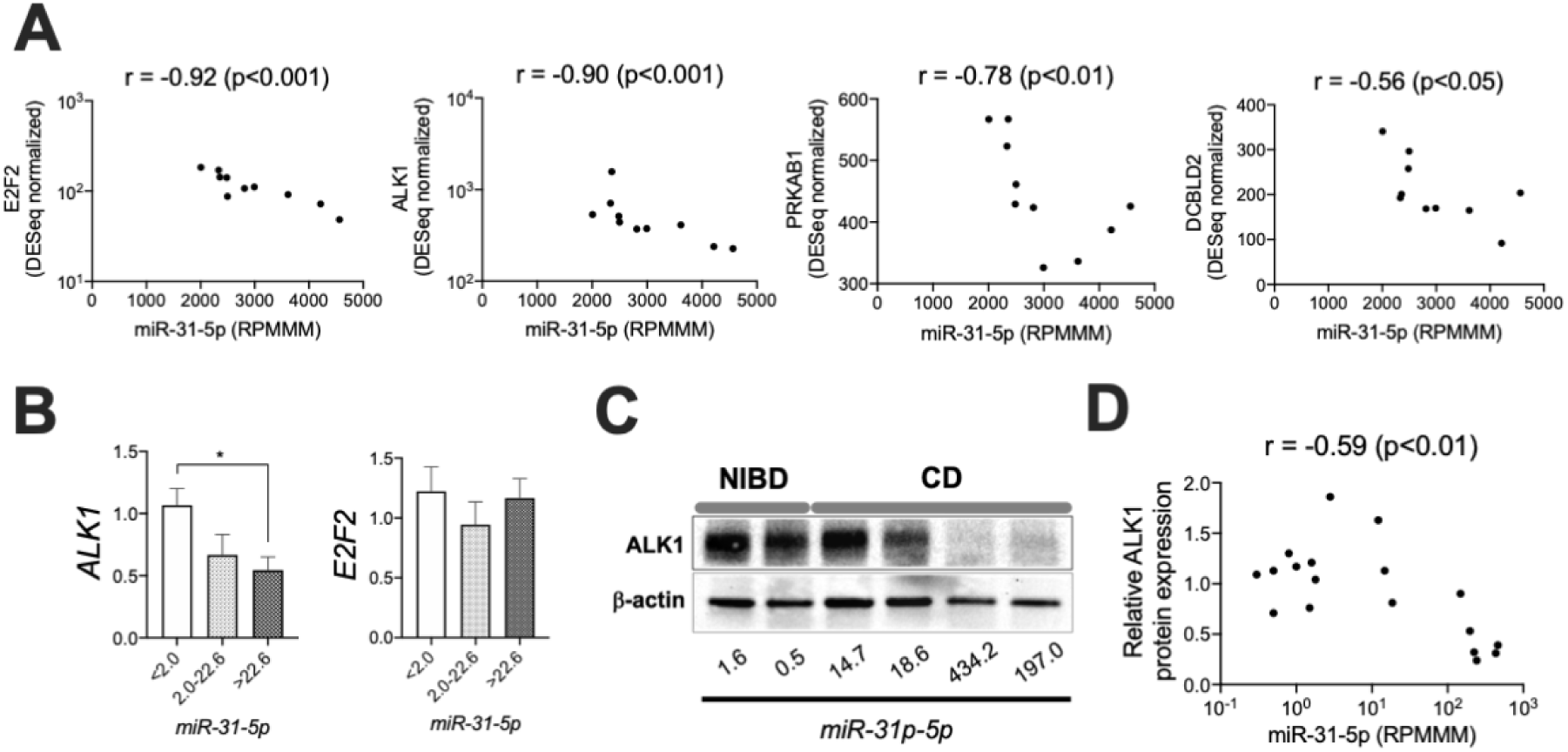
**(A)** Correlation of expression in the colonic mucosa of CD patients (N=10) between miR-31-5p (reads per million miRNAs mapped, RPMMM) and predicted targets of miR-31-5p (*E2F2, ALK1, PRKAB1*, and *DCBLD2*; DESeq normalized). **(B)** Association between the expression of miR-31-5p and the expression of *ALK1* or *E2F2* in isolated colonic epithelial cells (N=27). Gene expression was quantified by qPCR and samples were split into 3 equally sized groups (N=9 per group) according to the relative miR-31-5p expression levels. **(C)** Representative blot of ALK1 expression in the colonic tissue of NIBD and CD patients. (**D**) Correlation between ALK1 protein expression and miR-31-5p in the colonic mucosa (N=18). All correlation values were calculated by Spearman’s correlation coefficient. Each gene expression was normalized to *GAPDH* (*ALK1, E2F2*) or RNU48 (miR-31-5p). *P<0.05. P-values were determined by Kruskal-Wallis test followed by Dunn’s multiple comparison test.

### miR-31-5p directly suppress ALK1 expression through binding to the 3’UTR

To test the direct association between *ALK1* mRNA and miR-31-5p, we performed a 3’UTR reporter assay in HEK293T cells using synthesized miRNA and negative control mimics (**Figure 2A**). The mimics for miR-122a-5p and miR-215-5p, which do not have specific binding sites in the 3’UTR of *ALK1* mRNA, were used as negative controls. As we expected, miR-31-5p but not the other two miRNA mimics decreased luciferase activity compared to the mock and negative control mimic, suggesting the direct regulation of *ALK1* by miR-31-5p. To further demonstrate the specific regulation of *ALK1* by miR-31-5p through binding to the 3’UTR, we deleted the predicted miR-31-5p binding sequences in the reporter plasmid by site-directed mutagenesis (**Supplementary Figure 1**) and performed a reporter assay with miR-31-5p and negative control mimics (**Figure 2B**). The reduction in luciferase activity by exogeneous miR-31-5p mimic disappeared in the cells transfected with the mutated plasmid that does not have the intact miR-31-5p binding sequences. These results demonstrated the direct regulation of *ALK1* mRNA by miR-31-5p through binding to the 3’UTR.

**Fig. 2.**
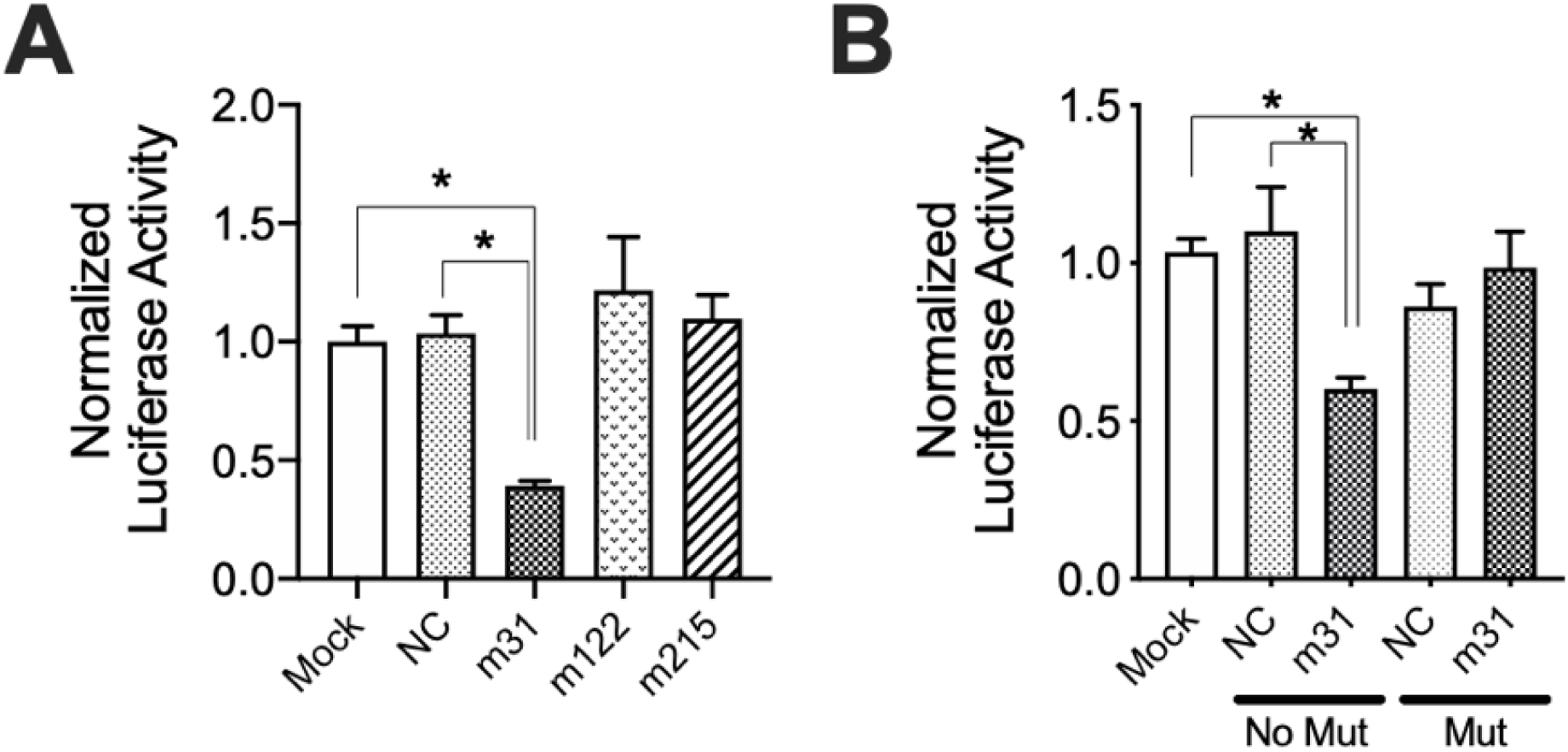
**(A)** 3’UTR reporter assay for *ALK1* in the presence or absence of 30nM miRNA mimics for hsa-miR-31-5p (m31), hsa-miR-122a-5p (m122), or hsa-miR-215-5p (m215) or negative control mimics (NC). **(B)** Site-directed mutagenesis assay with 10nM of m31 or NC mimics. *P<0.05. P-values were determined by Kruskal-Wallis test followed by Dunn’s multiple comparison test. N=6 per group for all experiments.

### BMP9-ALK1 signaling restricts the stemness of human colonic IECs

ALK1 is a type 1 receptor for transforming growth factor (TGF)-beta signaling molecules^24^ that specifically binds to extra-enteric ligands BMP9 and BMP10^25^. Although ALK1 is known to be expressed in human colonic IECs^26^, the role of ALK1-related signaling in IEC biology remains unknown. To investigate the impact of decreased ALK1 expression in the colonic IECs of CD patients, we first examined the effect of BMP9 treatment on IEC proliferation *ex vivo* using NIBD patient-derived colonic IEC monolayers. We found a significantly decreased EdU incorporation in BMP9-stimulated IECs compared to non-stimulated cells (**Figure 3A**). BMP9-induced reduction of EdU incorporation was recovered by sequestrating BMP9 with ALK1-Fc chimera protein, demonstrating the effect of BMP9 in restricting IEC proliferation. We also examined the effect of BMP9 on the expression of IEC proliferation- and stemness-related genes (**Figure 3B**). Consistent with the results of the EdU assay, the expression of proliferation-related genes, including *MKI67* and *PCNA* as well as stemness-related genes *LGR5, OLFM4*, and *ASCL2*, was significantly decreased in BMP9-stimulated cells. BMP9-induced reduction of expression levels of these genes was recovered by the addition of ALK1-Fc chimera protein.

**Fig. 3.**
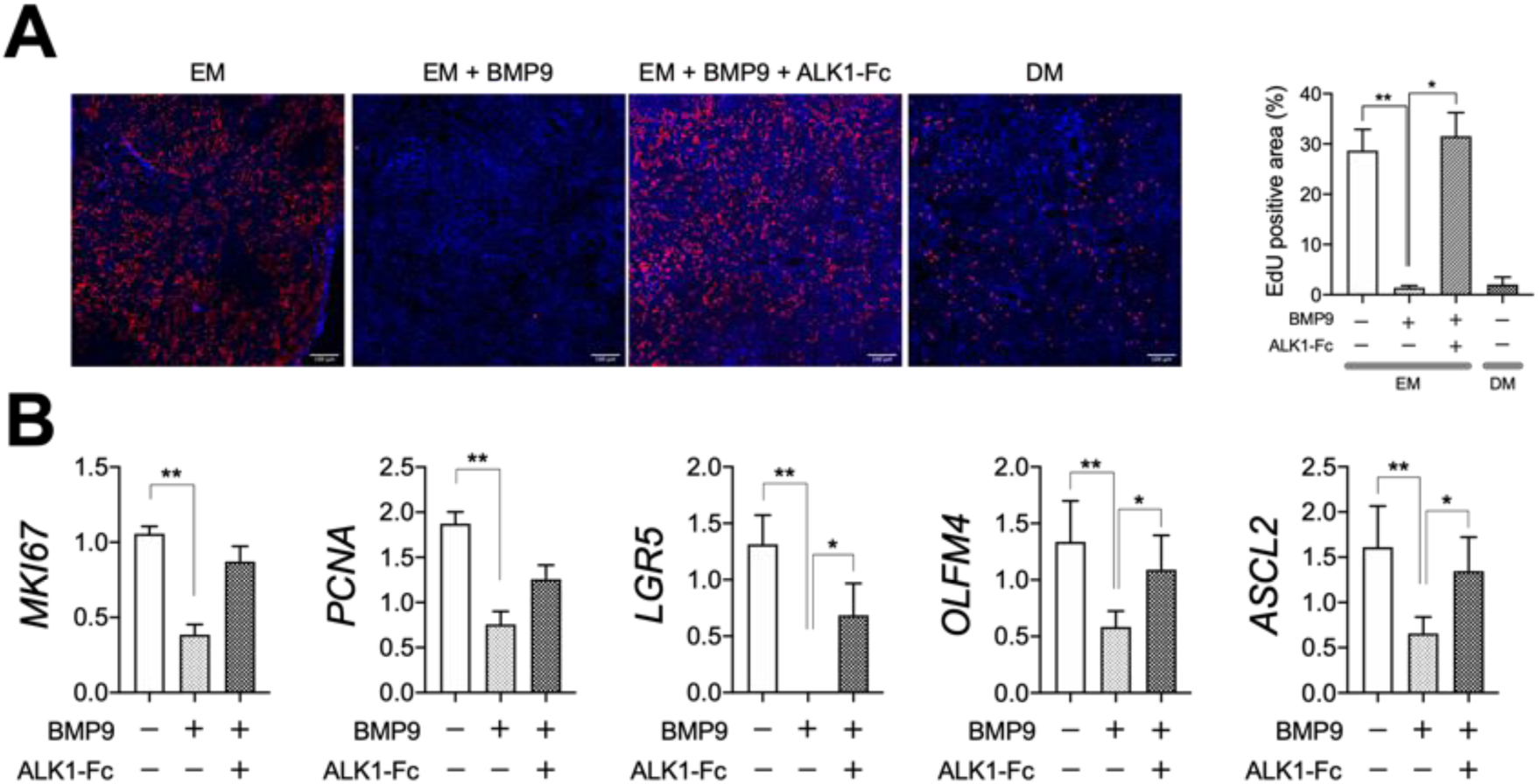
**(A)** EdU assay in NIBD patient-derived colonic epithelial cell monolayers (N=4-8 per group). Expanded cells were cultured in EM in the presence or absence of BMP9 and ALK1-Fc chimera protein. Red, EdU; Blue, Hoechst 33342. **(B)** Proliferation- and stemness-related gene expression in NIBD patient-derived colonic epithelial cell monolayers (N=6 per group). Each gene expression was normalized to *RPLP0*. *P<0.05, **P<0.01. P-values were determined by Kruskal-Wallis test (A) or Friedman test (B) followed by Dunn’s multiple comparison test. EM, expansion media; DM, differentiation media.

We hypothesized that colonic IECs of CD patients would maintain higher stemness than those of NIBD controls due to decreased ALK1 expression. To test this hypothesis, we compared proliferation- and stemness-related gene expression in the isolated colonic IECs between NIBD and CD patients (**Figure 4A**). As expected, CD patient-derived IECs exhibited significantly higher expression of *LGR5* and *OLFM4*, and a trend toward higher levels of *MKI67, PCNA*, and *ASCL2*, than those from NIBD controls. Increased OLFM4 protein expression was confirmed by immunohistochemistry in the colonic crypts of CD patients (**Figure 4B**). Taken together, these data suggest a role for BMP9-ALK1 signaling in restricting the stemness of human colonic IECs.

**Fig. 4.**
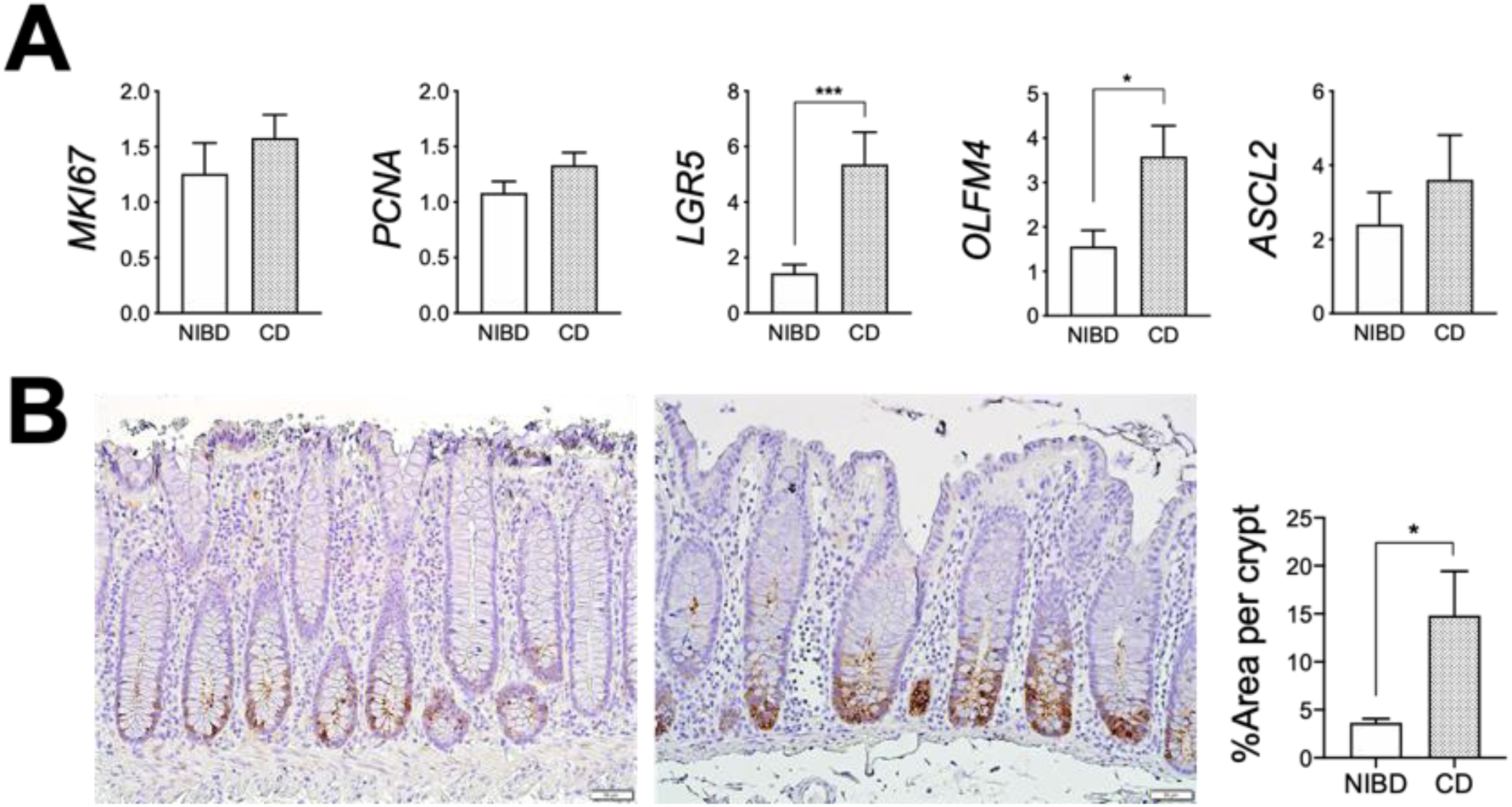
**(A)** Proliferation- and stemness-related gene expression in colonic epithelial cells isolated from CD patients (N=15) and NIBD controls (N=12). Each gene expression was normalized to *GAPDH*. (**B**) Representative immunohistochemical images of OLFM4 expression in the colonic crypts of NIBD controls (left) and CD patients (right). The percentage of OLFM4 staining area in colonic crypts was compared between CD patients and NIBD controls (N=4 per group). *P<0.05, ***P<0.001. P-values were determined by Mann-Whitney test.

### BMP9-ALK1 signaling drives colonic epithelial cell differentiation towards colonocytes

To define the impact of ALK1 signaling on IEC differentiation, we determined the expression of representative IEC-lineage specific genes^10, 27^, such as *CA1* (colonocyte), *MUC2* (goblet cell), *CHGA* (enteroendocrine cell), *DEFA5* (Paneth cell), and *DCLK1* (tuft cell), in NIBD patient-derived colonic IEC monolayers in the presence or absence of BMP9 stimulation. Treatment with BMP9 significantly upregulated the expression of *CA1* and other colonocyte markers^10^, such as *CEACAM1* and *GUCA2A* (**Figure 5A**), suggesting a potential impact of BMP9-ALK1 signaling on colonocyte differentiation. BMP9-mediated upregulation of *CA1* expression was confirmed at the protein level (**Figure 5B**). The pro-colonocytic effect of BMP9 was significantly mitigated by ALK1-Fc chimera protein (**Figure 5A**). These data demonstrate that BMP9-ALK1 signaling preferentially drives colonic IEC differentiation towards colonocytes.

**Fig. 5.**
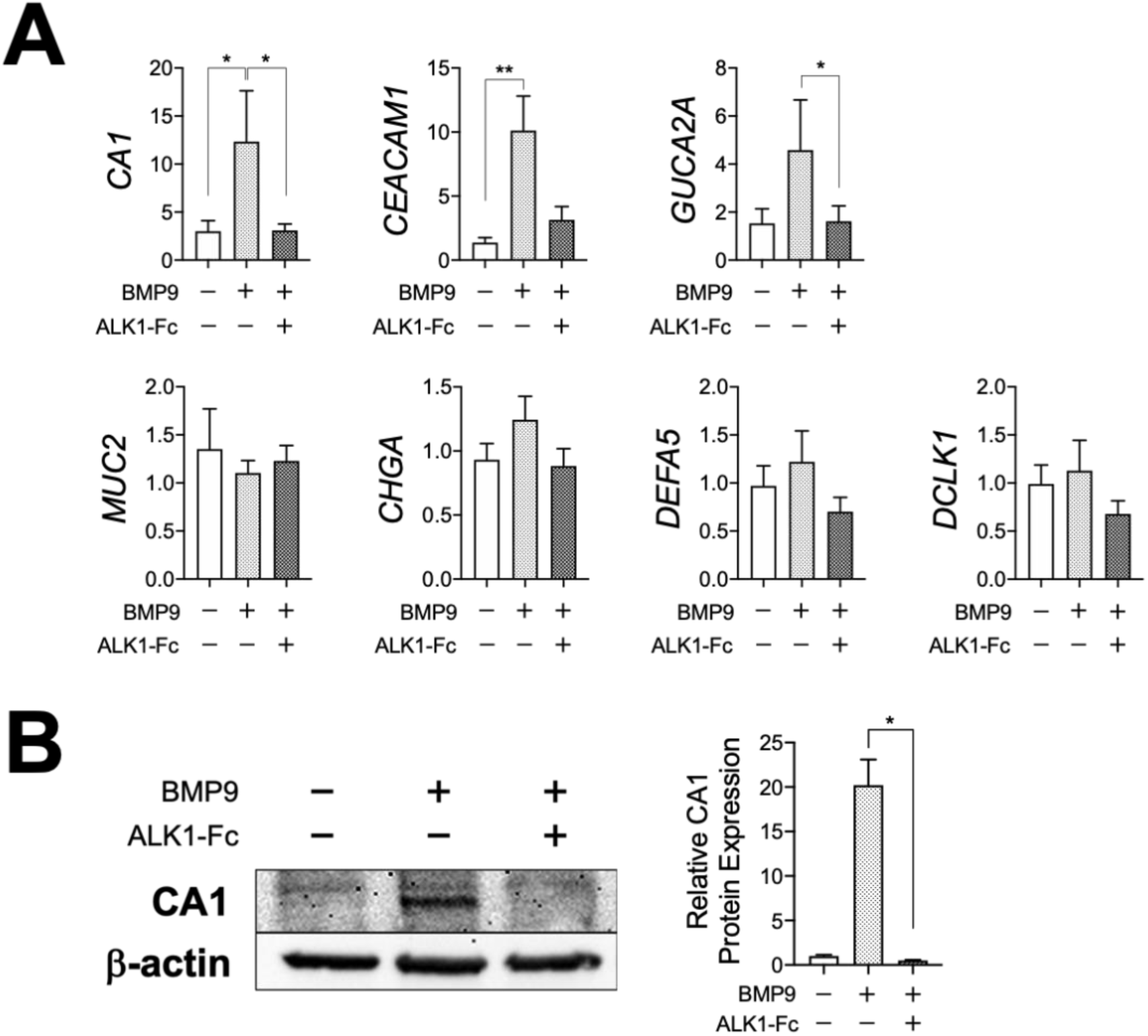
**(A)** Lineage-specific gene expression in NIBD patient-derived colonic epithelial cell monolayers. Expanded cells were cultured in expansion media in the presence or absence of BMP9 and ALK1-Fc chimera protein. Each gene expression was normalized to *RPLP0* (N=6 per group). **(B)** CA1 protein expression in NIBD patient-derived colonic epithelial monolayers (N=4 per group). *P<0.05, **P<0.01. P-values were determined by Friedman test followed by Dunn’s multiple comparison test.

We also hypothesized that decreased ALK1 expression in CD patients would result in decreased colonocyte marker expression compared to NIBD controls. To test this hypothesis, we compared expression of colonocyte markers in isolated colonic IECs between NIBD and CD patients (**Figure 6A**). CD patient-derived IECs showed significantly lower *CA1, CEACAM1*, and *GUCA2A* expression than IECs from NIBD controls. Reduced CA1 protein expression was also confirmed in the colonic mucosa of CD patients (**Figure 6B** and **6C**). Taken together, these data demonstrate for the first time the role of BMP9-ALK1 signaling in colonocyte differentiation.

**Fig. 6.**
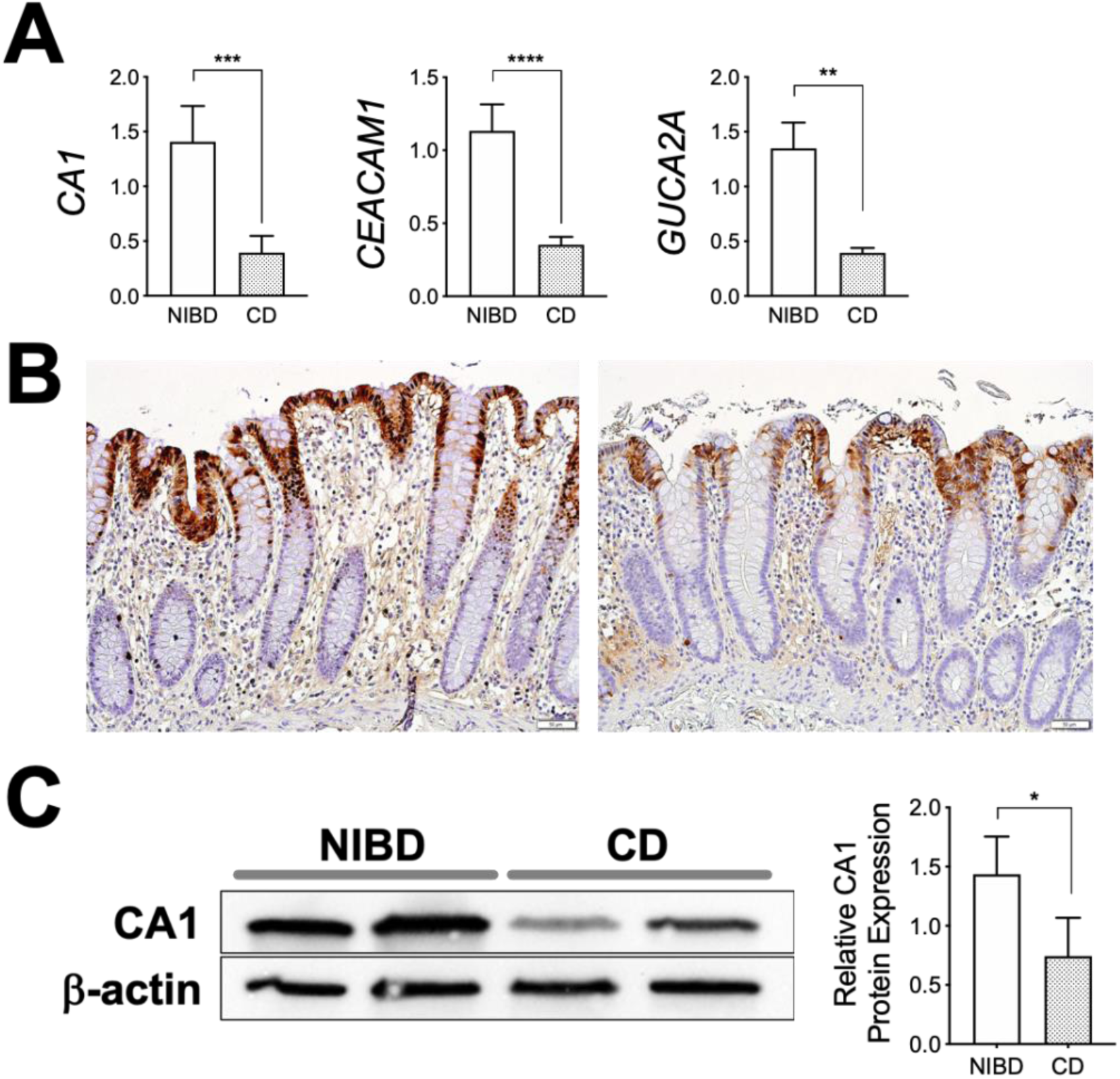
**(A)** Colonocyte marker expression in colonic epithelial cells isolated from CD patients (N=15) and NIBD controls (N=12). Each gene expression was normalized to *GAPDH*. (**B**) CA1 expression by immunohistochemistry in the colonic mucosa of NIBD controls (left) and CD patients (right). (**C**) Representative blot of CA1 protein expression in the colonic mucosa of NIBD controls and CD patients (N=5 per group in right bar graph). *P<0.05, **P<0.01, ***P<0.001, ****P<0.0001. P-values were determined by Mann-Whitney test.

### BMP9-ALK1 signaling does not affect migration of human colonic IECs

IECs migrate to wound sites and cover the denuded surface, an important step in the wound healing process^28^. We investigated the effect of BMP9-ALK1 signaling on wound healing using NIBD patient-derived colonic IEC monolayers. We measured how much of a wounded area was covered by migrated IECs within 24 hours after BMP9 stimulation (**Supplementary Figure 2**). The presence of BMP9 did not affect the amount of covered area at either 8 and 24 hours after stimulation. These results suggest that BMP9-ALK1 signaling has little or no impact on cell migration.

### BMP9-ALK1 signaling enhances human colonic IEC barrier integrity

Because colonocytes play a key role in barrier function, we hypothesized that ALK1 signaling could influence barrier integrity. To test this hypothesis, we generated IEC monolayers on transwells using NIBD patient-derived colonic crypts, which were cultured in the presence or absence of BMP9 (**Figure 7A**). BMP9 stimulation significantly enhanced IEC barrier integrity as shown by increased ΔTEER% (**Figure 7B**). This BMP9-induced increase in ΔTEER% was abrogated by ALK1-Fc chimera protein. These results suggest that increased BMP9-ALK1 signaling enhances human colonic IEC barrier integrity.

**Fig. 7.**
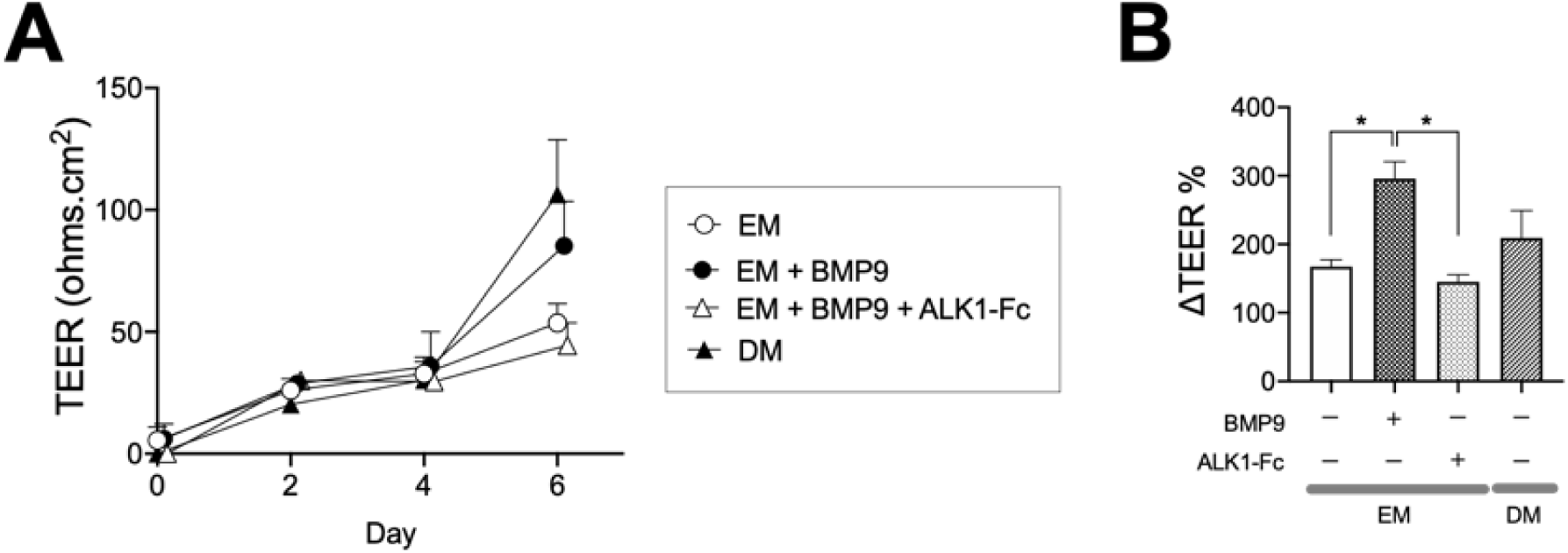
**(A)** Epithelial permeability assay in NIBD patient-derived colonic epithelial cells cultured on transwells. Trans-epithelial electrical resistance (TEER) was measured over time (N=3 per group). **(B)** Cells were stimulated with BMP9 in expansion media (EM) in the presence or absence of ALK1-Fc chimera protein or cultured in differentiation media (DM) at Day 4. The changes in TEER between Day 4 and Day 6 are shown as ΔTEER% (N=4-8 per group). *P<0.05. P-values determined by Kruskal-Wallis test followed by Dunn’s multiple comparison test.

### Decreased colonic ALK1 is associated with increased risk of surgery and endoscopic relapse in CD patients

We hypothesized that impaired ALK1 signaling, due in part to the associated defects in colonocyte maturation and barrier integrity, would be associated with poor clinical outcomes in CD. To study this, we examined *ALK1* expression in the colonic mucosa of a new, independent cohort of 28 CD patients and 10 NIBD controls using endoscopically-collected biopsy samples (**Figure 8A**). CD patients showed lower colonic *ALK1* expression than NIBD controls as a whole, but a subset of CD patients showed markedly lower *ALK1* expression (Low-ALK1 CD subset) than the other CD patients (Hi-ALK1 CD subset). Therefore, we compared the clinical characteristics between these ALK1-based CD subsets (**Table 1**). Interestingly, Low-ALK1 patients were more likely to experience bowel resections than Hi-ALK1 patients (93.3% vs. 53.8%, p=0.029). We examined the potential effect of surgery on colonic *ALK1* expression among 5 matched CD patients by prospectively determining *ALK1* expression after bowel resections. Within the median 6-month follow-up period, there was no significant change in the colonic *ALK1* expression (**Supplementary Figure 3A**). To evaluate the net impact of colonic ALK1 expression on the risk of surgery in CD patients, we conducted a multivariate analysis. In addition to well-accepted risk factors^29^ that include young age (under 40 years) at diagnosis, extensive disease (disease location^30^), use of corticosteroids to control the index flare, and perianal disease at diagnosis, we used membership in the colonic ALK1-based CD subset and use of anti-TNF alpha agents as additional independent variables that could impact the outcome (risk of surgery). Among these potential risk factors, young age at diagnosis was removed from subsequent analysis because of its significant correlation with membership in the colonic ALK1-based CD subset (**Supplementary Table 3** and **Supplementary Figure 3B**). A binomial logistic regression analysis revealed that having a Low-ALK1 CD subtype is an independent risk factor for surgical resection in CD patients (**Table 2**). Next, we prospectively tracked endoscopic disease activity in 21 patients who were in remission at the time of sample collection. Within a 72-week observational period, 5 of 12 Low-ALK1 and 0 of 9 Hi-ALK1 CD patients experienced endoscopic relapse. A significantly higher risk of endoscopic relapse was demonstrated in patients in the Low-ALK1 CD subset compared to the Hi-ALK1 subset (p<0.05, Log-rank test) (**Figure 8B**). All patients showed exacerbated inflammation in the colon except for one patient with endoscopic relapse only in ileum. Taken together, these data suggest that decreased colonic *ALK1* expression is associated with increased risk of surgery and endoscopic relapse in patients with CD.

**Table 1.** Clinical characteristics at the time of sample collection. P values were determined by ^a^Fisher’s exact test or ^b^Mann-Whitney test.

|  | Low-ALK1 | Hi-ALK1 | p value | Total |
| --- | --- | --- | --- | --- |
| total number | 15 | 13 |  | 28 |
| Gender, Male/Female, n (%) | 6 (40) / 9 (60) | 11 (84.6) / 2 (15.4) | 0.024<sup>a</sup> | 17 (60.7) / 11 (39.2) |
| Age at sampling, mean (SD) | 34.1 (8.8) | 40.4 (14.5) | 0.310<sup>b</sup> | 37.0 (12.0) |
| Disease duration (years), mean (SD) | 12.2 (9.1) | 11.6 (8.5) | 0.856<sup>b</sup> | 11.9 (8.7) |
| Former/Current smoker, n (%) | 7 (46.7) | 4 (30.8) | 0.460<sup>a</sup> | 11 (39.3) |
| Disease extent |  |  |  |  |
| L1/L2/L3, n (%) | 4 (26.7) / 1 (6.7) / 10 (66.7) | 4 (30.8) / 2 (15.4) / 7 (53.8) |  | 8 (28.6) / 3 (10.7) / 17 (60.7) |
| L4, n (%) | 3 (20) | 1 (7.7) | 0.600<sup>a</sup> | 4 (14.3) |
| Ileal involvement, n (%) | 14 (93.3) | 10 (76.9) | 0.311<sup>a</sup> | 24 (85.7) |
| Disease behaviour |  |  |  |  |
| B1/B2/B3, n (%) | 1 (6.7) / 11 (73.3) / 3 (20.0) | 4 (30.8) / 5 (38.5) / 4 (30.8) |  | 5 (17.9) / 16 (57.1) / 7 (25) |
| Perianal disease, n (%) | 5 (33.3) | 5 (38.5) | >0.999<sup>a</sup> | 10 (35.7) |
| Stricture, n (%) | 11 (73.3) | 5 (38.5) | 0.125<sup>a</sup> | 16 (57.1) |
| Disease activity |  |  |  |  |
| SES-CD, mean (SD) | 2.9 (3.1) | 4.4 (5.2) | 0.626<sup>b</sup> | 3.6 (4.2) |
| Endoscopic remission, n (%) | 12 (80.0) | 9 (69.2) | 0.670<sup>a</sup> | 22 (78.6) |
| Abnormal CRP ($\geq 5.0$mg/l), n (%) | 2/11 (18.2) | 0/8 (0.0) | 0.485<sup>a</sup> | 2/19 (10.5) |
| Treatment history |  |  |  |  |
| Aminosalicylates, n (%) | 0 (0.0) | 3 (23.1) | 0.087<sup>a</sup> | 3 (10.7) |
| Oral steroids, n (%) | 4 (26.7) | 5 (38.5) | 0.689<sup>a</sup> | 9 (32.1) |
| Immunosuppressants, n (%) | 7 (46.7) | 4 (30.8) | 0.460<sup>a</sup> | 11 (39.3) |
| Anti-TNF alpha agents, n (%) | 9 (60.0) | 6 (46.2) | 0.705<sup>a</sup> | 15 (53.6) |
| Surgery, n (%) | 14 (93.3) | 7 (53.8) | 0.029<sup>a</sup> | 21 (75.0) |

**Table 2.** Binomial logistic regression analysis for surgery. Standard Deviation, SD; odds ratio, OR; confidential interval, CI.

|  | Regression Coefficient | SD | p value | OR | 95% CI of OR |
| --- | --- | --- | --- | --- | --- |
| Low-ALK1 CD subset | 2.91 | 1.37 | 0.03 | 18.33 | 1.26 to 266.39 |
| Disease location | 0.85 | 0.68 | 0.21 | 2.34 | 0.62 to 8.81 |
| Corticosteroids | -0.38 | 1.46 | 0.80 | 0.68 | 0.04 to 11.93 |
| Anti-TNF alpha agents | -0.86 | 1.17 | 0.46 | 0.42 | 0.04 to 4.21 |
| Perianal disease | 0.61 | 1.30 | 0.64 | 1.85 | 0.14 to 23.63 |

**Fig. 8.**
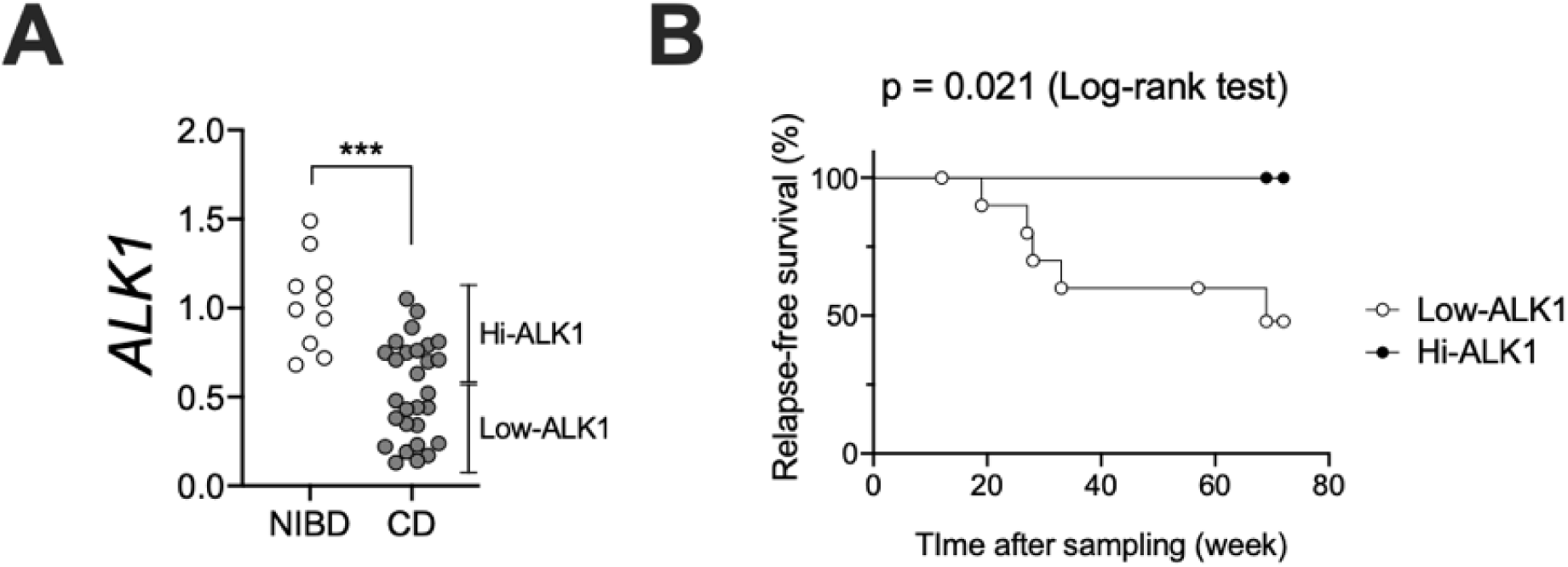
**(A)** The impact of colonic *ALK1* expression on clinical outcome. *ALK1* expression was quantified in colonic biopsy samples obtained from NIBD controls (N=10) and CD patients (N=28) by qPCR. *ALK1* expression was normalized to *GAPDH*. ***P<0.001. P-value was determined by Mann-Whitney test. **(B)** Kaplan-Meier survival analysis to evaluate the impact of colonic ALK1 expression on the endoscopic relapse in patients with CD (N=12 for Low-ALK1 and 9 for Hi-ALK1 CD subgroups). P-value determined by Log-rank test.

## Discussion

ALK1 is a transmembrane serine/threonine receptor kinase that belongs to the TGF-beta receptor family. In contrast to the broad ligand-binding specificity of other type 1 receptors for TGF-beta signaling molecules (ALK2-7), ALK1 specifically binds to BMP9 and BMP10 secreted from liver^31^ and heart^32^, respectively. ALK1 is expressed in endothelial cells and plays a crucial role in the development of direct connections between arteries and veins^33^.

Defects in ALK1 signaling lead to arteriovenous malformations and cause hereditary hemorrhagic telangiectasia (HHT), which is characterized by mucocutaneous telangiectasia and gastrointestinal hemorrhage, in genetically predisposed patients^34^. Recent tissue proteome analyses revealed ALK1 expression in human IECs as well as endothelial cells in small and large intestines^26^. However, the role of ALK1 signaling in colonic IEC homeostasis and the clinical impact of attenuated ALK1 signaling in CD have never been investigated. In this manuscript, we identified *ALK1* as a target of miR-31-5p^4, 5^. We demonstrated a dramatic reduction of ALK1 in the colonic IECs of CD patients and showed that ALK1 controls colonocyte differentiation and intestinal epithelial barrier capacity.

BMP4 signaling through ALK3 was previously reported to restrict the stemness of murine Lgr5-positive intestinal stem cells^35^. Although ALK1 and ALK3 similarly affect downstream Smad signaling pathways to regulate gene expression^24^, the role of ALK1-related signaling on IEC stemness remained unknown. Therefore, we examined the impact of ALK1 signaling on the stemness of human colonic IECs *ex vivo* using primary-cultured IECs. BMP9-ALK1 signaling reduced the expression of stemness-related genes including *LGR5*, restricted proliferation, and directed the fate of human IEC differentiation toward colonocytes. These data clearly demonstrated the effect of BMP9-ALK1 signaling in restricting human IEC stemness, similar to BMP4-ALK3 signaling. Decreased ALK1 expression was associated with increased stemness-related marker expression and decreased colonocyte differentiation in the colonic IECs of CD patients compared to NIBD controls. We also demonstrated that increased BMP9-ALK1 signaling enhanced IEC barrier integrity by decreasing paracellular permeability. Although the mechanism of decreased paracellular permeability in BMP9-stimulated IECs needs further investigation, the effect of BMP9-ALK1 signaling in colonocyte differentiation and the role of colonocytes in colonic barrier integrity^9^ implies that activated ALK1 signaling might enhance barrier integrity by driving colonocyte differentiation and maturation.

Based on these data, we hypothesized that attenuated ALK1 signaling might result in a poor clinical outcome in CD patients due to disrupted colonic barrier function. Association analyses with clinical outcomes revealed that decreased colonic *ALK1* expression is an independent risk factor for surgery and is associated with an increased risk of endoscopic relapse in the future. Despite the demonstrated role of ALK1 signaling in epithelial barrier function, loss-of-function mutations in ALK1 do not result in spontaneous intestinal inflammation in patients with HHT^36^. This fact in consistent with previous lines of evidence from rodents^37-39^ and asymptomatic first-degree relatives of CD patients^2^ suggesting that increased epithelial permeability is necessary but not sufficient for the development of intestinal inflammation and requires another trigger for disease manifestation in CD^40^. Nonetheless, our data suggested that disrupted ALK1 signaling might be one of the disease-modifying factors that contribute to the heterogeneity in clinical outcomes among CD patients. Finding markers predicting the natural course of CD is the holy grail of medical researchers in the field^41^. Our findings suggest that colonic ALK1 expression levels may help identify these high-risk patients who may require early intensified therapeutic intervention, perhaps staving off a poor clinical outcome. Intriguingly, we found a significant correlation between colonic ALK1-based CD subset and young age at diagnosis as independent variables for the regression analysis. The younger age at diagnosis in Low-ALK1 CD subset compared to Hi-ALK1 subset might partially explain the unknown mechanism why young age at diagnosis can be a risk factor for the disabling course in CD patients. Our findings also show a potential use of ALK1 as a novel target of medical treatment in CD. An increasing number of drugs targeting mucosal immune cells are currently available in the treatment for CD; however, there is no established treatment targeting IEC defects^8^. A small biologic potentiator of colonic ALK1 expression could be a first-ever treatment targeting IECs in CD.

There are several limitations in our study. Firstly, we did not investigate the impact of decreased E2F2, another candidate target of miR-31-5p, in CD patients. E2F2 is a transcriptional factor that plays an important role in regulating cell proliferation^42, 43^. Our data suggested that miR-31-5p regulation of E2F2 expression does not occur in colonic IECs of CD patients. However, increased miR-31-5p was also found in CD14 negative resident macrophages and B lymphocytes in the colon of CD patients^4^. Future investigation will be required to understand the potential impact of decreased E2F2 expression in these non-epithelial cells. Secondly, we did not examine the impact of ALK1 signaling on IECs from the small intestine. We previously reported an upregulation of miR-31-5p in the ileal tissue of CD patients compared to NIBD controls^4^. Given that ALK1 is expressed in the IECs of small intestine as well as colon^26^, investigation of the role of ALK1 signaling in small intestinal IECs will be informative. Thirdly, we did not investigate the impact of decreased ALK1 on vascular development in the intestine. ALK1 has a well-accepted role in vascular development, thus increased miR-31-5p might affect vascular formation in the intestine of CD patients. Finally, we did not examine the effect of BMP10, another ligand for ALK1, on ALK1 signaling in IECs. Both BMP9 and BMP10 can bind to ALK1 with high affinity *in vivo*. However, in contrast to BMP9 which circulates in a biologically active form^31^, there remains an active debate about the latency of BMP10^44^. Therefore, we selected BMP9 for our *ex vivo* experiments. Nevertheless, distinct roles of these two BMPs were previously reported in tumor growth, and thus further investigation of the impact of BMP10 on IEC biology will be necessary in the future.

To our knowledge, this study is the first to investigate the mechanism of CD using human primary IEC monolayers *ex vivo*. Our innovative technologies, such as cell proliferation, wound healing, and epithelial permeability assays in 2D-cultured primary IECs, will promote further research to understand mechanisms of IBD and develop novel drugs targeting IECs in the future.

## Supplementary methods

### Isolation of Colonic Epithelial Cells

Colonic epithelial cells were isolated as previously described^4^. In brief, dissected colonic mucosa was cut into small pieces and incubated in magnesium-free HBSS containing 2mM EDTA and 2.5% heat-inactivated fetal bovine serum (FBS) for 30 minutes with shaking at 37°C. To remove mucus, 1 mM dithiothreitol was added and incubated for additional 10 minutes with shaking. Collected supernatants were centrifuged, re-suspended in HBSS containing 1 mg/ml collagenase type 3 (Worthington Biochemical, NJ, USA), and incubated for 10 minutes at 37°C to further remove the mucus. The fraction was pelleted, re-suspended in HBSS, passed through 40um filter, and overlayered on 50% Percoll. Cells were centrifuged at 2000 rpm for 20 minutes at room temperature and viable colonic IECs were recovered from the interface layer.

### Quantitative Reverse-Transcriptase Polymerase Chain Reaction Analysis (qPCR)

Total RNA was extracted from dissected colonic mucosa using TRIzol Reagent and purified with Total RNA Purification Kit (Norgen Biotek, ON, Canada) according to the manufacturer’s instruction. Total RNA was extracted from isolated and cultured colonic IECs using Single Cell RNA Purification Kit (Norgen Biotek). cDNA for mRNA was generated from 500ng of RNA using High-Capacity cDNA Reverse Transcription Kit (Thermo Fisher Science, MA, USA). cDNA for miRNAs was generated from 10ng of RNA using TaqMan MicroRNA Reverse Transcription Kit (Thermo Fisher Science) with individual miRNA-specific primers (TaqMan miRNA Assays, Assay ID: 001006 [RNU48], 002279 [miR-31-5p]). qPCR for mRNAs was performed on QuantStudio 3 RT-PCR system using PowerUp SYBR Green Master Mix (Thermo Fisher Science) or BrightGreen 2x qPCR MasterMix-Lox ROX (Applied Biological Materials, BC, Canada). Each gene expression was normalized to Glyceraldehyde 3-phosphate dehydrogenase (*GAPDH*) or Ribosomal Protein Lateral Stalk Subunit P0 (*RPLP0*). qPCR for miRNAs was performed using TaqMan Universal PCR Master Mix with individual miRNA-specific probes (Thermo Fisher Science). Expression of *miR-31-5p* was normalized to *RNU48*.

### Immunohistochemistry

Immunostaining was performed as described previously^45^. Rabbit anti-human CA1 antibody, Rabbit anti-human OLFM4 antibody, and Mouse/Rabbit IgG VisUCyte HRP polymer antibody were purchased from Novus Biologicals (CO, USA), Cell Signaling Technology (MA, USA), and R&D Systems. The images were captured on an inverted microscope (Olympus IX71, Olympus, Tokyo, Japan) with cellSens Standard software.

### Western Blot Analysis

Western blot analyses were performed on whole-cell extracts^45^. Goat anti-human ALK1 antibody and Goat IgG HRP-conjugated antibody were purchased from R&D Systems. Rabbit anti-human CA1 and β-actin antibodies were obtained from Novus Biologicals and Cell Signaling Technology, respectively. The image was captured on a chemiluminescent image reader (iBright Imaging Systems, Invitrogen, MA, USA) and analyzed by ImageJ software. Expression of ALK1 and CA1 protein expression was normalized to β-actin.

### Wound healing assay

Colonic IECs were expanded on 12-well cell culture inserts (BD Falcon) coated with collagen hydrogel for 4 days to make confluent IEC monolayers. After cultured in DM for 2 days, 1 mm^2^ of cells were cut off from 8 different portions of monolayers by biopsy punches to generate cell free “wounded” areas. To stop cell proliferation and examine the net impact of BMP9 stimulation on cell migration, cells were subsequently cultured in DM lacking FBS in the presence or absence of 10 ng/ml BMP9 for an additional 24 hours. The wounded areas were imaged with a confocal microscope at 8 and 24 hours. The average percentages of the initial wounded areas covered by migrating cells were evaluated using ImageJ software.

---

**Supplementary Fig 1.**
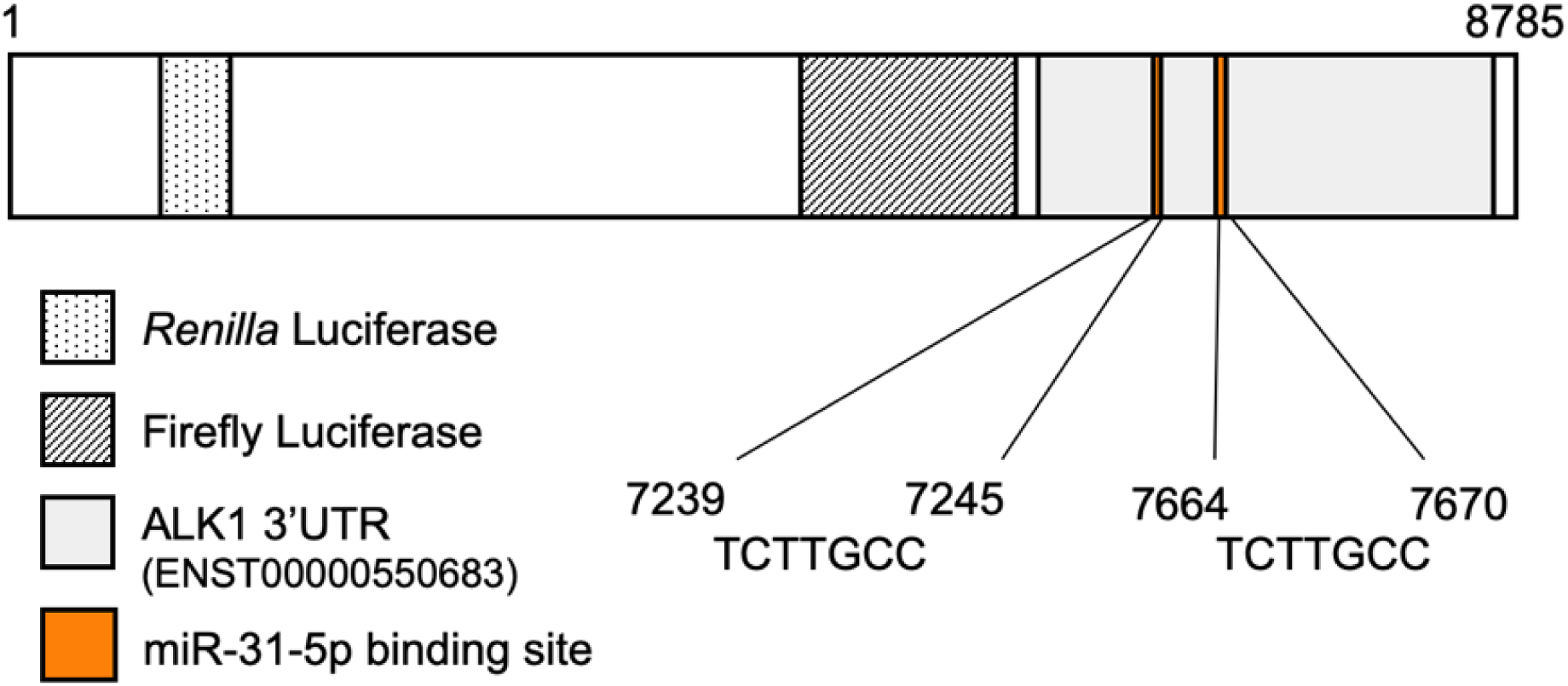
Schematic representation of the miR-31-5p binding sites in the reporter plasmid.

**Supplementary Fig 2.**
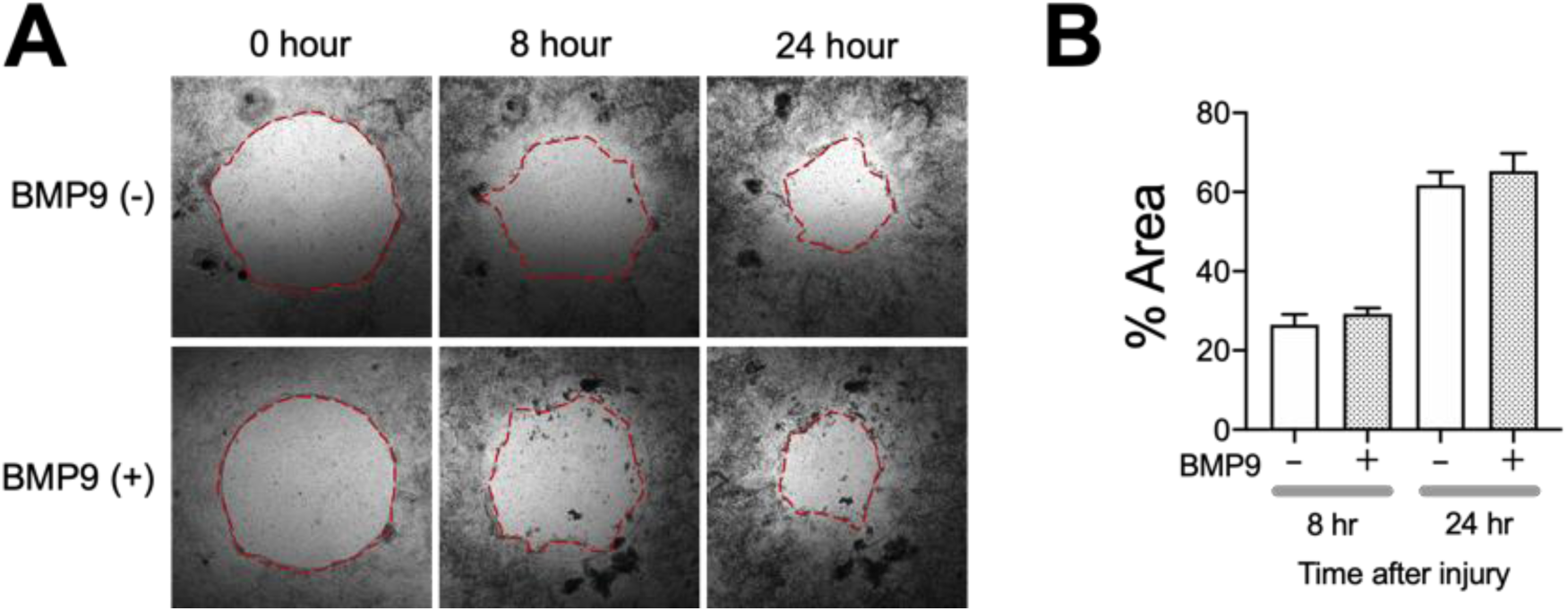
**(A)** Wound healing assay in NIBD patient-derived colonic epithelial cell monolayers. **(B)** The wounded area covered with migrated cells were measured at 8 and 24 hours (hr) after BMP9 stimulation (N=7 per group).

**Supplementary Fig 3.**
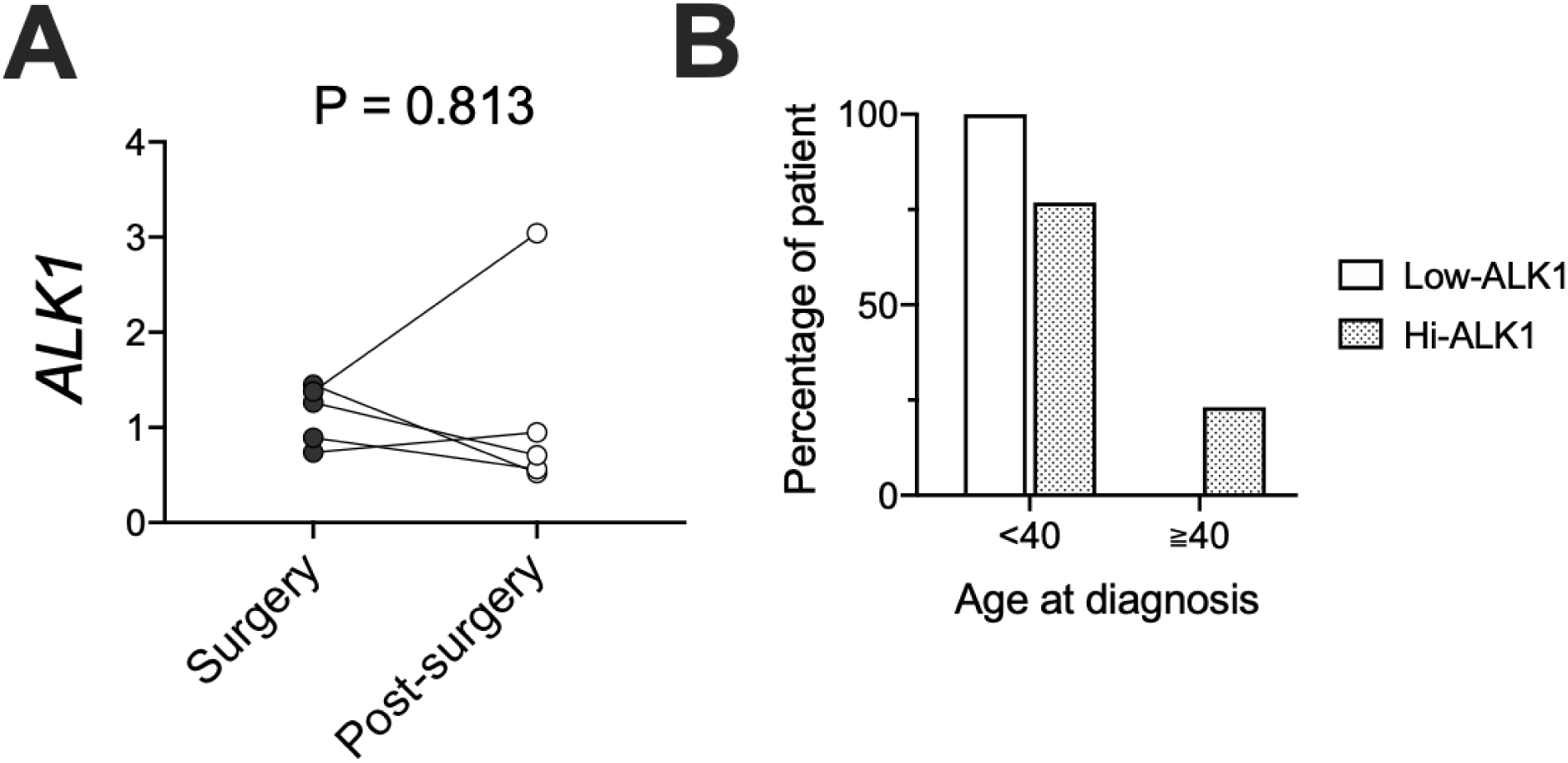
**(A)** *ALK1* expression in the colonic mucosa of CD patients at the time of surgery and after surgery (N=5 per group). *ALK1* expression was normalized to *GAPDH*. P-value was determined by Wilcoxon test. **(B)** Percentages of CD patients who were diagnosed as CD before and after 40 years old in Low-ALK1 (N=15) and Hi-ALK1 (N=13) CD subsets.

**Supplementary Table 1.** Clinical information of surgical samples. Right side, cecum ∼ ascending colon; Transverse, transverse colon; Left side, descending colon ∼ sigmoid colon.

| Age (year) | Gender | Disease | Sample Location |
| --- | --- | --- | --- |
| 68 | Male | Adenoma/Benign tumor | Left side |
| 62 | Male | Adenoma/Benign tumor | Right side |
| 74 | Female | Adenoma/Benign tumor | Right side |
| 52 | Female | Colon cancer | Right side |
| 51 | Male | Colon cancer | Transverse |
| 80 | Female | Colon cancer | Right side |
| 43 | Male | Colon cancer | Right side |
| 50 | Male | Colon cancer | Left side |
| 83 | Female | Colon cancer | Right side |
| 44 | Male | Colon cancer | Left side |
| 44 | Female | Colon Inertia | Left side |
| 41 | Female | Colon Inertia | Right side |
| 51 | Female | Constipation | Right side |
| 50 | Female | Crohn's disease | Right side |
| 66 | Female | Crohn's disease | Right side |
| 44 | Male | Crohn's disease | Rectum |
| 21 | Male | Crohn's disease | Left side |
| 13 | Male | Crohn's disease | Right side |
| 49 | Female | Crohn's disease | Right side |
| 28 | Female | Crohn's disease | Right side |
| 20 | Male | Crohn's disease | Right side |
| 45 | Female | Crohn's disease | Right side |
| 18 | Male | Crohn's disease | Right side |
| 49 | Female | Crohn's disease | Right side |
| 47 | Male | Crohn's disease | Right side |
| 29 | Female | Crohn's disease | Right side |
| 26 | Male | Crohn's disease | Right side |
| 75 | Male | Crohn's disease | Right side |
| 26 | Male | Crohn's disease | Right side |
| 45 | Female | Crohn's disease | Transverse |
| 53 | Female | Crohn's disease | Right side |
| 35 | Male | Crohn's disease | Right side |
| 44 | Female | Crohn's disease | Right side |
| 43 | Female | Crohn's disease | Right side |
| 34 | Male | Crohn's disease | Right side |
| 16 | Female | Crohn's disease | Right side |
| 52 | Male | Crohn's disease | Rectum |
| 53 | Female | Crohn's disease | Right side |
| 33 | Male | Crohn's disease | Right side |
| 23 | Male | Crohn's disease | Right side |
| 18 | Male | Crohn's disease | Right side |
| 23 | Male | Crohn's disease | Right side |
| 70 | Male | Crohn's disease | Right side |
| 42 | Male | Crohn's disease | Right side |
| 45 | Female | Crohn's disease | Right side |
| 27 | Female | Crohn's disease | Right side |
| 68 | Male | Diverticulitis | Left side |
| 64 | Female | Diverticulitis | Left side |
| 73 | Female | Diverticulitis | Left side |
| 58 | Female | Diverticulitis | Left side |
| 54 | Male | Diverticulitis | Left side |
| 48 | Female | Diverticulitis | Left side |
| 76 | Male | Diverticulitis | Left side |
| 54 | Female | Diverticulitis | Left side |
| 47 | Male | Diverticulitis | Left side |
| 64 | Female | Diverticulitis | Left side |
| 65 | Female | Diverticulitis | Left side |
| 60 | Male | Ischemic colitis | Right side |
| 76 | Female | Sigmoid volvulus | Left side |
| 49 | Female | Submucosal tumor | Right side |
| 64 | Male | Submucosal tumor | Right side |

**Supplementary Table 2.** Twenty-seven predicted miR-31-5p target genes which were significantly down-regulated in colonic mucosa of high-miR-31-5p CD patients compared to NIBD controls.

| Gene symbol | Average (DESeq normalized count) |  | log2FoldChange | Adjusted p value |
| --- | --- | --- | --- | --- |
|  | NIBD | high-miR31 CD |  |  |
| AKAP7 | 290.5 | 141.1 | -1.00 | 0.00015 |
| ALK1 | 1298.3 | 538.9 | -1.19 | 0.00027 |
| AK4 | 667.7 | 261.5 | -1.24 | 0.00075 |
| ARRDC3 | 1211.1 | 771.1 | -0.63 | 0.00625 |
| ATP10B | 3285.4 | 2057.8 | -0.65 | 0.01031 |
| COL4A4 | 99.2 | 60.7 | -0.72 | 0.00628 |
| DCBLD2 | 396.0 | 208.8 | -0.88 | 0.00266 |
| E2F2 | 297.2 | 115.5 | -1.29 | <0.00001 |
| FAM118B | 385.7 | 244.7 | -0.65 | 0.00007 |
| FMN1 | 1240.0 | 670.1 | -0.86 | 0.00023 |
| HOMER1 | 98.5 | 35.2 | -1.29 | 0.00420 |
| KIAA1462 | 509.3 | 152.0 | -1.29 | 0.04777 |
| KPNA4 | 1808.1 | 1183.4 | -0.59 | 0.00973 |
| MATR3 | 434.8 | 204.5 | -1.02 | 0.00136 |
| MBOAT2 | 1019.3 | 390.1 | -1.32 | <0.00001 |
| NARS2 | 246.7 | 168.8 | -0.52 | 0.01588 |
| ORC5 | 165.4 | 128.0 | -0.36 | 0.03969 |
| PPP1R12B | 3463.1 | 1188.0 | -1.33 | 0.00589 |
| PPP1R9A | 607.3 | 291.7 | -1.04 | 0.00002 |
| PRKAB1 | 577.6 | 444.7 | -0.37 | 0.04906 |
| RAB6B | 71.5 | 33.9 | -0.94 | 0.03396 |
| SATB2 | 2296.5 | 468.0 | -2.22 | <0.00001 |
| SLC26A2 | 22136.2 | 2596.6 | -2.86 | <0.00001 |
| TLN2 | 1492.2 | 755.4 | -0.95 | 0.00004 |
| VAV3 | 1141.7 | 522.4 | -1.07 | 0.00013 |
| ZC3H12C | 894.9 | 463.3 | -0.91 | 0.00011 |
| ZNF704 | 1412.6 | 820.8 | -0.77 | 0.00001 |

**Supplementary Table 3.** Correlations between explanatory variables for the risk of surgery. Correlations between categorical variables were analyzed by Chi-squared test. *p<0.05. The strength of correlations measured by Cramer’s V test was shown in the table.

| **Correlations** | ALK1 subset | Age<40 at diagnosis | Disease location | Corticosteroids | anti-TNF alpha | Perianal disease |
| --- | --- | --- | --- | --- | --- | --- |
| ALK1 subset |  | 0.372* | 0.161 | 0.045 | 0.138 | 0.053 |
| Age<40 at diagnosis | 0.372* |  | 0.270 | 0.292 | 0.141 | 0.017 |
| Disease location | 0.161 | 0.270 |  | 0.339 | 0.324 | 0.243 |
| Corticosteroids | 0.045 | 0.292 | 0.339 |  | 0.045 | 0.141 |
| anti-TNF alpha | 0.138 | 0.141 | 0.324 | 0.045 |  | 0.096 |
| Perianal disease | 0.053 | 0.017 | 0.243 | 0.141 | 0.096 |  |

